# Impact of Fluorophore and Epitope Position on Destabilized Reporter Performance in *C. elegans*

**DOI:** 10.64898/2026.04.10.717803

**Authors:** Anton Jackson, James Matthew Ragle, Jordan D. Ward

## Abstract

Determining spatial and temporal gene expression is crucial for understanding animal development and physiology. Promoter reporters are powerful tools used to dissect how cis-regulatory elements and trans-acting factors control gene expression. Many fluorescent proteins used in promoter reporters, however, have long half-lives (>24 hr) which limit the study of dynamic expression. Destabilizing sequences like PEST reduce the half-life of reporter proteins to provide a more representative readout of gene expression. *mlt-11* is a putative protease inhibitor known to oscillate in expression at the mRNA and protein level, yet a *mlt-11p::mNeonGreen::3xFLAG::PEST::tbb-2 3’UTR* promoter reporter did not detectably oscillate. We systematically dissected this transgene, finding that placement of a 3xFLAG tag adjacent to a PEST sequence severely hampered oscillatory expression of a mNeonGreen promoter reporter. Surprisingly, reporter designs that effectively oscillate with a GFP or mNeonGreen fluorophore fail to oscillate when mStayGold is used, with these reporters remaining detectable over 24 hours following promoter inactivation. In addition, other tested epitopes (Myc, ALFA, OLLAS) did not hamper PEST-dependent destabilization but led to varying levels of reporter expression. This study details key considerations for designing destabilized fluorescent promoter reporters.

## INTRODUCTION

Dynamic gene expression underlies many key biological processes, such as biological timers, cell differentiation, stress and immune responses, neural activity, and metabolic state transitions (Sheng and Greenberg 1990; West and Greenberg 2011; Ramirez et al. 2017; Semkova et al. 2022; Zeng et al. 2024; Spangler et al. 2025; Zhao et al. 2025). Promoter reporters are a powerful tool to monitor promoter activity in living animals and assay the role of cis-regulatory elements and trans-acting factors in the regulation of a given gene. These tools involve taking the promoter of a gene of interest and fusing it with a reporter gene that produces a detectable protein such as LacZ, luciferase, or a fluorescent protein (FP)(Casadaban 1976; de Wet et al. 1987; Chalfie et al. 1994). One limitation in using FP-based promoter reporters is that FPs often have long half-lives. For example, green fluorescent protein (GFP) has a half-life of ∼26 hours (Corish and Tyler-Smith 1999). This stability hampers study of dynamic promoter activity, as persisting reporter protein can mask promoter inactivation. To counter this limitation, many researchers fuse a destabilizing sequence to their reporter, reducing reporter protein half-life to two hours (Li et al. 1998). The most commonly used sequence is a proline, glutamic acid, serine, and threonine-rich “PEST” sequence from mouse ornithine decarboxylase (Loetscher et al. 1991). It promotes protein degradation through a ubiquitin-independent proteasomal pathway, mediated by the antizyme protein, with portable activity that destabilizes the nuclear or cytosolic proteins to which it is fused (Murakami et al. 1992; Zhang et al. 2003).

*mlt-11* is a Kunitz domain-containing putative protease inhibitor required to pattern all layers of the cuticle and to promote embryonic viability in *C. elegans* (Ragle et al. 2026). Both its mRNA and protein have been demonstrated to oscillate, peaking in the middle of each larval stage (Frand et al. 2005; Hendriks et al. 2014; Meeuse et al. 2020; Ragle et al. 2026). Curiously, we observed that a *mlt-11p::mNeonGreen::3xFLAG::PEST::tbb-2 3’UTR* promoter reporter generated for that study (Ragle et al. 2026) was constitutively expressed (Fig. 1). Here, we systematically dissect this reporter, revealing that the 3xFLAG tag suppressed PEST-induced destabilization. GFP and mNeonGreen (mNG) were destabilized by a PEST sequence when the 3xFLAG tag was either omitted or placed on the opposite terminus of mNG relative to the PEST sequence. Surprisingly, mStayGold (mSG), a bright, photostable, green FP, resists PEST-induced destabilization and remains detectable over 24 hours after promoter activity ceases. The inactivation function of the 3xFLAG tag is unique to this epitope, as 3xMyc, 3xOLLAS, and 3xALFA tags allow PEST to function, though they impact reporter intensity to variable levels. Together these studies provide guidelines for promoter reporter design, depending on whether brightness or dynamics are prioritized, and for epitope and PEST placement for destabilized reporters.

**Figure 1.**
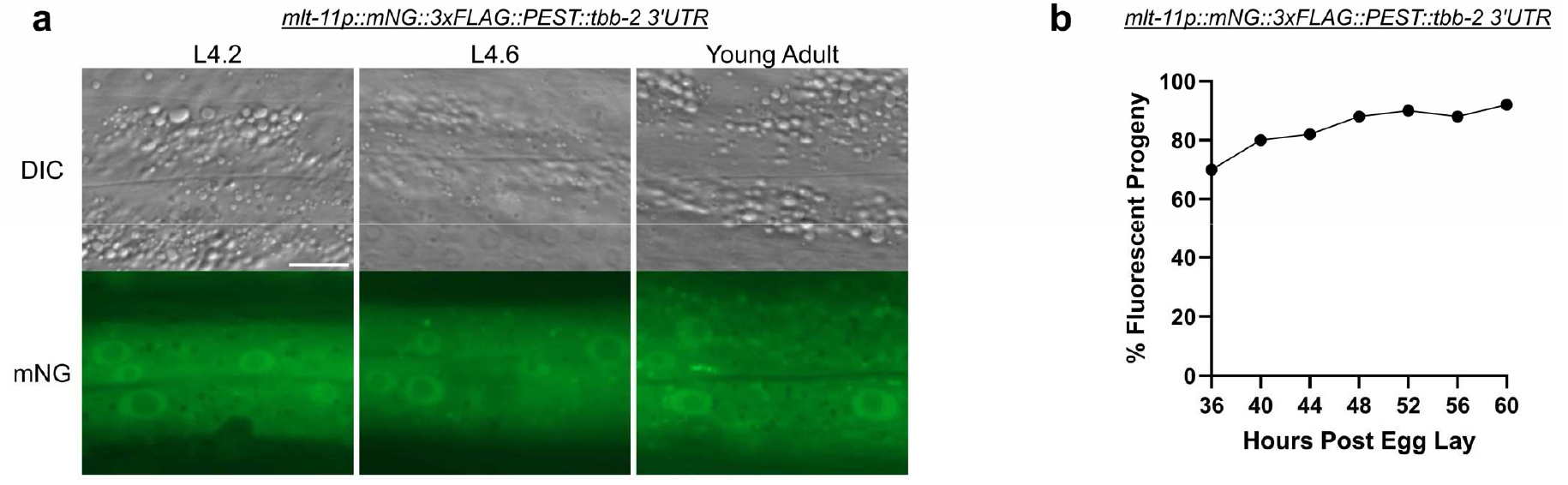
*mlt-11p::mNG::3xFLAG::PEST* reporters fail to oscillate. (a) Larval stage 4.2 (L4.2), larval stage 4.6 (L4.6), and young adult images of a *mlt-11p::mNG::3xFLAG::PEST::tbb-2 3’UTR* promoter reporter. Scale bar=10 µm. Images are representative of 20 animals scored over two independent experiments. (b) Scoring activity of a *mlt-11p::mNG::3xFLAG::PEST::tbb-2 3’UTR* promoter reporter over a 24 hour period. The fluorescent microscope LED light intensity was set to 33% during the course of the experiment. 50 animals were scored over two independent experiments.

## RESULTS

### GFP and mNeonGreen reporters oscillate when fused with a PEST Sequence

A *mlt-11p::mNG::3xFLAG::PEST::tbb-2 3’UTR* promoter reporter failed to recapitulate previously reported oscillatory mRNA and protein expression (Fig. 1)(Frand et al. 2005; Hendriks et al. 2014; Meeuse et al. 2020; Ragle et al. 2026). As this reporter was dim and challenging to screen, we shifted to nuclear-localized *mlt-11p::GFP::H2B::unc-54 3’UTR* fusions, imaging three developmental timepoints which capture a pulse of *mlt-11* expression (L4 stage 4.6) flanked by periods of low expression (L4 stage 4.2, young adulthood). This reporter used the same promoter fragment that was shown to oscillate in Frand et al., 2005. As most constructs in this study use a standard *unc-54* 3’ UTR for somatic cell expression, all reporters will use this 3’UTR unless otherwise noted. In addition, we synchronized animals via a timed egg lay and scored promoter reporter containing animals over a 24 period under a fluorescent dissecting microscope. We expected to detect three pulses of *mlt-11* expression and tested two different LED light intensities to determine whether that impacted reporter detection. We validated key observations using the oscillatory *qua-1* promoter (Meeuse et al., 2020).

A GFP control reporter lacking the PEST sequence and driven by either *mlt-11p* or *qua-1p* failed to oscillate as expected (Fig 2a to c), whereas GFP reporters containing an N-terminal PEST sequence oscillated (Fig. 2d to f), recapitulating data from previous studies (Frand et al. 2005; Meeuse et al. 2020). We observed the same oscillatory pattern when scoring at 100% or 33% light intensity. We then tested how mNG and mSG performed as reporters, using the same two promoters. The PEST::mNG::H2B construct performed similarly to a GFP equivalent (Fig. 2g to i), indicating that a PEST sequence can destabilize an mNG::H2B reporter. Surprisingly, PEST::mSG::H2B did not detectably oscillate (Fig. 2j to l). The PEST::mSG::H2B reporter follows similar expression patterns to that of GFP::H2B, with little to no oscillation (Fig. 2j to l).

**Figure 2.**
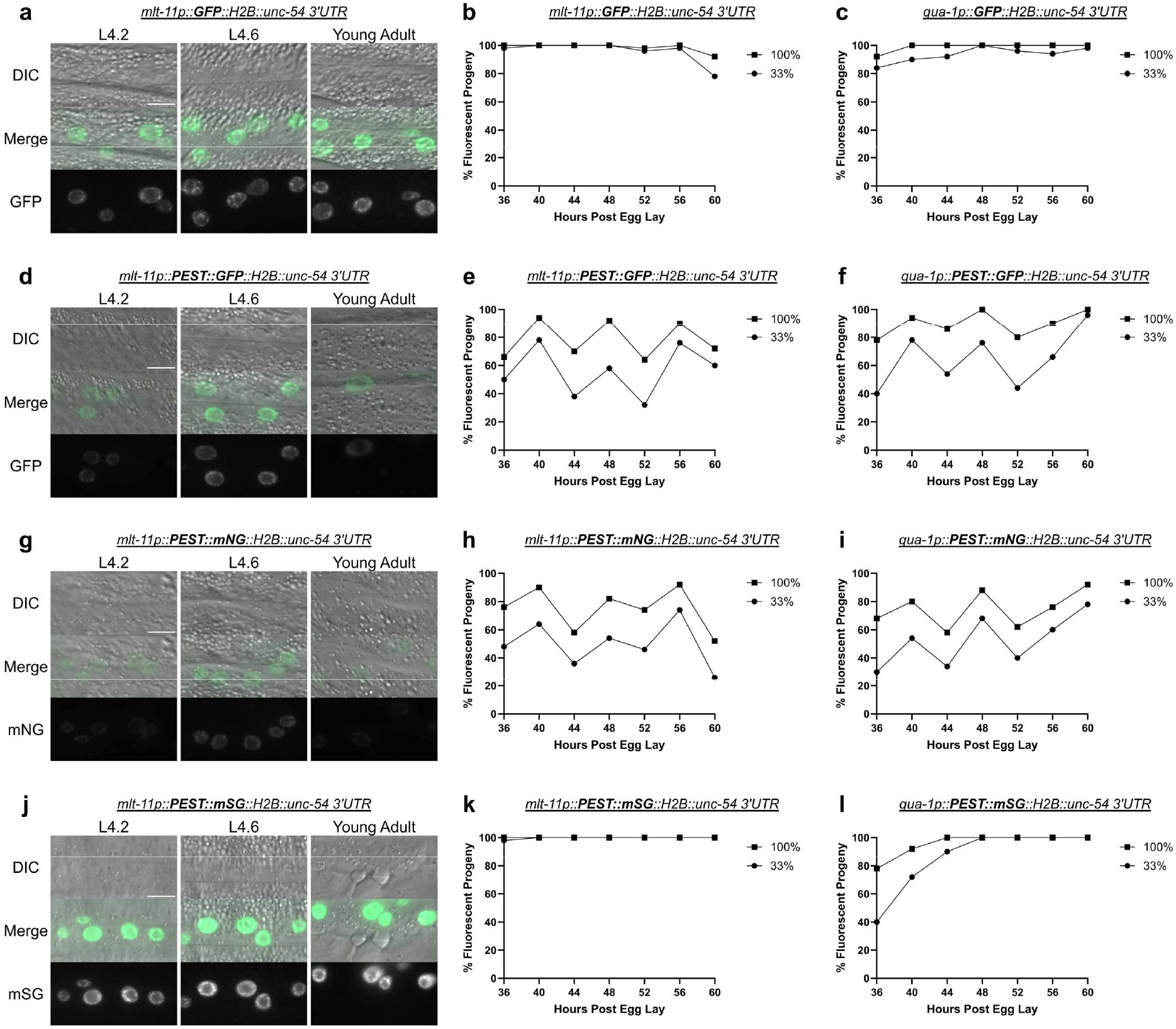
Comparison of PEST destabilization function on GFP, mNG, and mSG reporters. Representative images of GFP::H2B (a), PEST::GFP::H2B (d), PEST::mNG::H2B (g), and PEST::mSG::H2B (j) driven by *mlt-11p* at the indicated stages (Larval stage 4.2 (L4.2), larval stage 4.6 (L4.6), and young adult). Scale bars=10µm. Scoring expression of a *GFP::H2B::unc-54 3’UTR* promoter reporter driven by *mlt-11p* (c) and *qua-1p* (d). Scoring expression of *PEST::GFP::H2B::unc-54 3’UTR* promoter reporters driven by *mlt-11p* (e) and *qua-1p* (f). Scoring expression of *PEST::mNG::H2B::unc-54 3’UTR* promoter reporters driven by *mlt-11p* (h) and *qua-1p* (i). Scoring expression of *PEST::mSG::H2B::unc-54 3’UTR* promoter reporters driven by *mlt-11p* (k) and *qua-1p* (l). Animals for the time course experiments were scored over a 24 hour period with the LED light intensity set to 100% or 33% as noted in the figure legend. Two independent experiments were performed; images are representative of 20 animals and expression was scored in 50 animals.

### mSG reporters are bright and PEST sequences do not destabilize them

In our comparative experiments (Fig. 2) we noted that PEST::mSG::H2B is significantly brighter than GFP and mNG equivalents (Fig. 2d, g, e, Fig 3a and b). The GFP and mNG promoter reporters were not significantly different in brightness while mSG was 5.25 and 5.36 times brighter than GFP and mNG respectively (Fig. 3a). Given that the PEST sequence did not appear to effectively destabilize the mSG reporter, we extended our time course to determine how long it took reporter activity to decline. The *mlt-11* promoter becomes inactive in mid-to-late L4 and the gene is not detectably expressed in adults (Frand et al. 2005; Meeuse et al. 2020; Ragle et al. 2026). PEST::GFP::H2B declined rapidly in adulthood (64-68 hours post-egg lay; Fig 3c). In contrast, the PEST::mSG::H2B reporter was fully on at 72 hours post-egg lay, and then slowly declined in expression with ∼40% of animals still expressing detectable reporter at 84 hours post-egg lay when scored under 100% light intensity (Fig. 3c).

**Figure 3.**
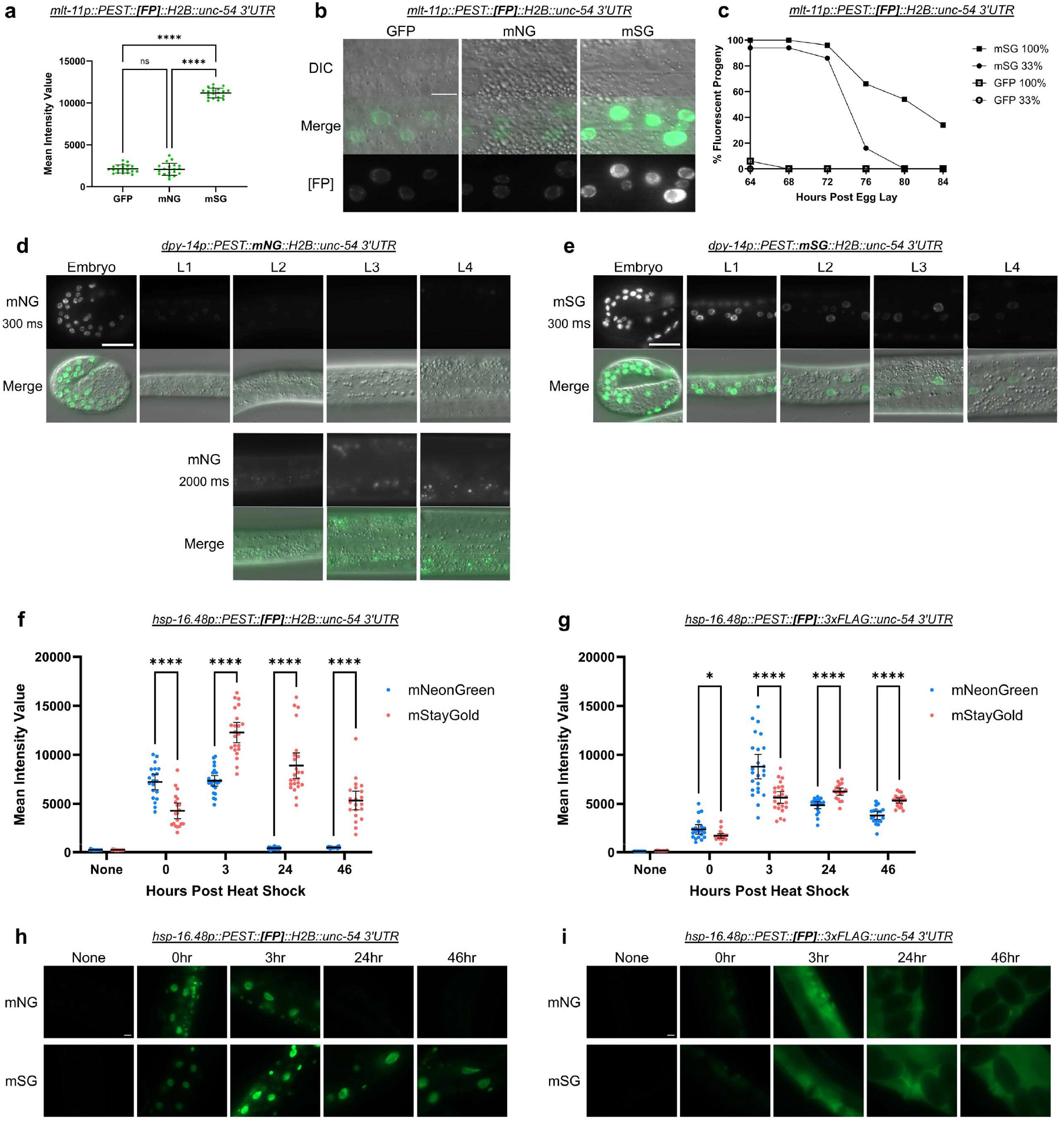
mStayGold is not destabilized by a PEST sequence: (a) Mean intensity value of three promoter reporters 48 hours post-timed egg lay. A one-way ANOVA with Tukey’s multiple comparisons test was performed to determine significance. (b) Representative images of the promoter reporters quantified in Fig. 3a. Scale bar=10µm. 20 worms were imaged over two experiments in a and b. (c) Scoring expression of the indicated promoter reporters. Fifty animals from two independent experiments were scored over a 24-hour period starting 64-hour post egg-lay at the indicated LED light intensities. (d and e) Representative images of the indicated promoter reporters at stages ranging from embryogenesis to L4 with either 300 ms (top panel d, e) or 2000 ms (bottom panel d) exposure. (f and g) Mean intensity values of the indicated promoter reporters before a 34°C heat-shock, and after at the indicated timepoints. (h and i) Representative images of the promoter reporters quantified in Fig. 3f and 3 g, respectively. To determine significance, a Welch’s T-test was performed between each fluorescent protein at the indicated hours in Fig. 3f and g. ns=p≥0.05, *=p<0.05, ****=p<0.0001. Scale bar=20 µm for d, e, h, and i. Images are representative of 50 animals scored over two independent experiments in d, e, h, and i. Data from 15-25 animals imaged over two independent experiments were used for the quantitation in f and g.

These results suggested that PEST::mSG::H2B was not destabilized by the PEST sequence. To further test this finding, we generated *dpy-14p::PEST::[FP]::H2B* promoter reporters (Fig. 3d to e). The *dpy-14* gene is exclusively expressed during *C. elegans* embryogenesis (Gallo et al. 2006). If PEST::mSG::H2B was unable to be degraded, we predicted that there would be continued expression in the L2 stage and beyond. A control *dpy-14p::PEST::mNG::H2B* strain showed strong fluorescence in embryos, with expression dimming at the L1 stage (Fig. 3d). Increasing the exposure time from 300 ms to 2000 ms showed that at the L2 stage and beyond, there was little to no detectable mNG expression. In contrast, in the *dpy-14p::PEST::mSG::H2B* strain, reporter expression persists into the L4 stage despite the promoter becoming inactive as animals hatch to L1 larvae (Fig 3e)(Gallo et al. 2006; Gerstein et al. 2010). Together, these data indicate that, in an H2B reporter, mSG is significantly brighter than mNG and GFP, and PEST sequences fail to destabilize mSG.

As this study had been focused on promoters predominantly expressed in the epidermis, we compared PEST::mNG and PEST::mSG reporter expression driven by a heat shock promoter, with and without an H2B fusion. For nuclear-localized H2B reporters, we saw higher PEST::mNG::H2B expression when animals were imaged immediately after a one hour heat shock (0 hr; Fig 3f and h). Both reporters peaked in expression by 3 hours post-heat shock, with PEST::mSG::H2B exhibiting higher expression. The PEST sequence destabilized the mNG::H2B reporter as we observed low reporter activity after 24 hours post-heat shock, while the PEST::mSG::H2B was still robustly expressed 46 hours post-heat shock (Fig. 3f and h). In contrast, reporters lacking the H2B displayed different expression dynamics. Weak PEST::mNG::3xFLAG and PEST::mSG::3xFLAG were detected at the 0 hour timepoint. At the 3 hour timepoint both reporters were robustly expressed with PEST::mSG::3xFLAG displaying comparatively weaker expression (Fig. 3g and i). Reporter activity slowly declined, with PEST::mSG::3xFLAG animals being brighter both 24 and 46 hours post-heat shock and both reporters remaining detectable at the 46 hour timepoint (Fig. 3g and i). This work further confirms that PEST does not destabilize mSG and surprisingly indicated that PEST seems far more effective at destabilizing an mNG::H2B reporter compared to an unlocalized reporter.

### A 3xFLAG tag suppresses reporter oscillation when adjacent to a PEST sequence

As mNG::H2B was destabilized by a PEST sequence, we then explored why our initial *mlt-11p::mNG::3xFLAG::PEST::tbb-2 3’ UTR* reporter failed to oscillate (Fig. 1). The 3xFLAG tag was an obvious candidate to test. We generated a comparable H2B reporter (*mlt-11p::mNG::3xFLAG::PEST::H2B)* to the Fig. 1 reporter and it failed to oscillate (Fig. 4a and 4b). In all cases, we separated 3xFLAG and PEST with a flexible 11 amino acid linker. Similar effects were seen with an equivalent GFP reporter (Fig. 4c). We next explored whether the position of the PEST sequence relative to the 3xFLAG epitope affected this suppression of destabilization function. PEST was moved to the N-terminus of the reporter, which restored oscillatory expression (Fig. 4d and e). We validated this result using a *qua-1* promoter reporter, which also showed oscillation when scored (Fig. 4f). Finally, similar effects were observed in an equivalent *mlt-11p::PEST::GFP::3xFLAG::H2B* reporter (Fig. 4g).

**Figure 4.**
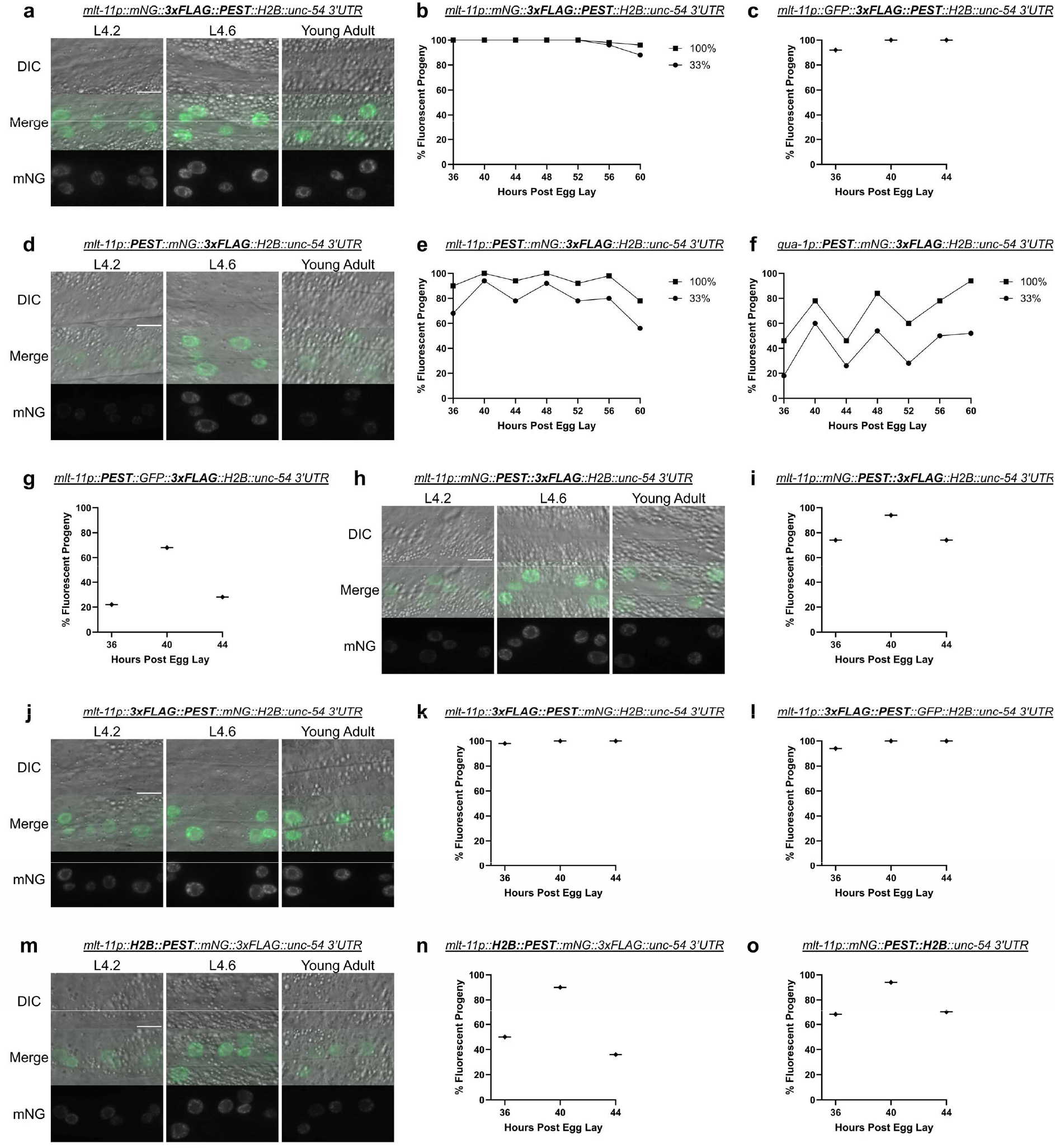
3xFLAG N-terminal to PEST sequences prevents destabilization of GFP::H2B and mNG::H2B reporters: Representative images of mNG::3xFLAG::PEST::H2B (a), PEST::mNG::3xFLAG::H2B (d), mNG::PEST::3xFLAG::H2B (h), 3xFLAG::PEST::mNG::H2B (j), and H2B::PEST::mNG::3xFLAG (m) all driven by *mlt-11p* at the indicated stages (Larval stage 4.2 (L4.2), larval stage 4.6 (L4.6), and young adult). Scale bars=10 µm. (b, c, e, f). Scoring expression of the indicated promoter reporters. Animals for the time course experiments were scored over a 24 hour period starting 36 hours post egg-lay with the LED light intensity set to 100% or 33% as noted in the figure legend. (g, i, k, l, n, o) Scoring expression of the indicated promoter reporters over a 8 hour period (36-44 hr) post-egg lay with the LED light intensity set to 10%. Two independent experiments were performed; images are representative of 20 animals and expression was scored in 50 animals.

We next tested whether the inactivation function of the 3xFLAG tag was specific to being N-terminal to the PEST sequence. A *mlt-11p::mNG::PEST::3xFLAG::H2B* reporter exhibited a modest oscillation (Fig. 4h and i) compared to equivalent reporters with the 3xFLAG being N-terminal to PEST (Fig. 4a and b). 3xFLAG::PEST on the N-terminus of the fluorophore also hampered oscillation in both mNG and GFP reporters (Fig. 4j to l). Given the effect of 3xFLAG on PEST function, two reporters were designed to test whether the H2B could affect PEST-induced destabilization. An *H2B::PEST::mNG::3xFLAG* reporter oscillated, though there appeared to be higher reporter activity in the expression troughs in L4.2 and young adulthood (Fig. 4m and n). A *mlt-11p::mNG::PEST::H2B* reporter showed dampened oscillation though not as severe as the effect of 3xFLAG N-terminal to PEST (Fig. 4k, 4o). Together, these data indicate that flanking sequences can affect PEST function with N-terminal 3xFLAG having the most potent disruption to PEST activity, and 3xFLAG or H2B C-terminal to the PEST sequence having more moderate effects.

### Other common epitopes vary in reporter expression

We next tested whether 3xMyc, 3xALFA, 3xOLLAS, and 3xHA epitopes had similar inhibitory effects on PEST function (Evan 1985; Field 1988; Park 2010; Götzke 2019). We generated a set of *mlt-11p::[epitope]::PEST::mNG::H2B* reporters, with the epitopes separated from the PEST sequence by the same 11 amino acid flexible linker used for the 3xFLAG reporters, and monitored their expression (Fig. 5). 3xMyc, 3xALFA, and 3xOLLAS permitted oscillation (Fig. 5a to c). Surprisingly, the 3xHA reporter showed little to no expression at any of the developmental stages analyzed (Fig. 5d). This data may suggest that HA enhances PEST activity or somehow impacts gene expression. In addition, from L4.6 to young adult, there was a statistically significant decrease in expression levels for all promoter reporters except in the case of 3xHA (Fig. 5e). While most epitopes permitted oscillation, the level of expression significantly varied among reporters (Fig. 5f). 3xMyc reporters had the largest mean intensity value at the L4.6 stage, being approximately 1.4 times brighter than 3xALFA, 2.1 times brighter than 3xOLLAS, and 14.7 times brighter than 3xHA. These results indicate that 3xMyc, 3xALFA, and 3xOLLAS minimally impact reporter destabilization when adjacent to PEST while 3xHA severely dampens reporter expression when on the N-terminus adjacent to PEST.

**Figure 5.**
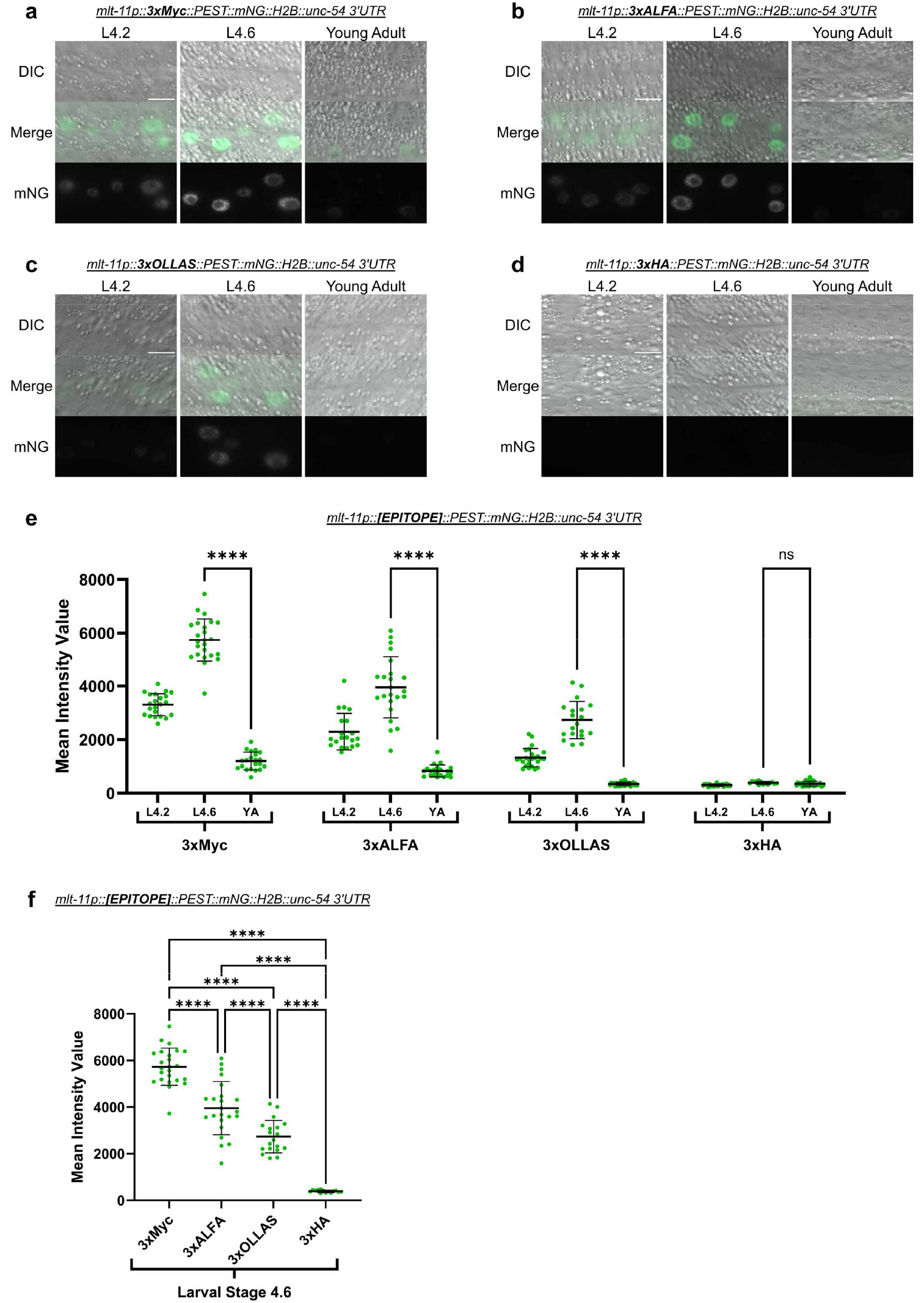
Comparison of epitope effect on PEST destabilization function. (a to d) Representative images of the indicated *mlt-11p::[epitope]::PEST::mNG::H2B::unc-54 3’ UTR* promoter reporters at larval stage 4.2 (L4.2), 4.6 (L4.6), and young adulthood. Scale bars=10 µm. (e) Measured fluorescence intensity of the indicated genotypes at the L4.2, L4.6, and young adult timepoints. Images of 14-27 animals per strain were collected over two experiments and their average fluorescence was used to generate mean intensity value data. A Welch’s t-test was performed between each strain’s L4.6 and young adult (YA) means to determine significance. (f) Comparison of the L4.6 stages from the data collected in Fig 5e. A one-way ANOVA with Tukey’s multiple comparisons test was performed to determine significance between the L4.6 group of each epitope. ns=p≥0.05, ****=p<0.0001.

## DISCUSSION

Based on this work, we recommend that researchers using PEST-based promoter reporters consider (1) the fluorescent protein being used and (2) the presence of epitopes next to the PEST sequence itself. PEST was sufficient to degrade mNG::H2B and GFP::H2B reporters. PEST::mSG::H2B reporters failed to oscillate and exhibited persistent expression. The presence of an N-terminal 3xFLAG tag potently suppressed PEST-dependent destabilization, with oscillatory expression being restored by separating the two tags. Other common epitopes generally supported oscillation when adjacent to PEST, but varied in reporter expression levels.

### PEST sequences fail to destabilize mSG reporters

This study began when a *mlt-11p::mNG::3xFLAG::PEST::tbb-2 3’UTR* promoter reporter showed no oscillation despite the *mlt-11* mRNA and protein oscillating during *C. elegans* development (Fig. 1)(Frand et al. 2005; Hendriks et al. 2014; Meeuse et al. 2020; Ragle et al. 2026). We initially focused on whether the issue might be the fluorophore. Control GFP::H2B reporters that lacked (Fig. 2a to c) or contained PEST (Fig. 2d to f) were used to compare how mNG and mSG reporters behaved. PEST::mNG::H2B reporters (Fig. 2g to i) performed similarly to PEST::GFP::H2B equivalents, indicating that PEST can effectively destabilize mNG::H2B. Interestingly, PEST::mSG::H2B reporters (Fig. 2j to l) failed to oscillate, similar to the GFP::H2B control reporter. PEST::mSG::H2B was significantly brighter than the PEST::GFP::H2B and PEST::mNG::H2B equivalents (Fig. 3a and b). In addition, PEST::mSG::H2B was still detectable well into adulthood unlike a PEST::GFP::H2B reporter (Fig 3c). A *dpy-14p::PEST::mSG::H2B* promoter reporter remained detectable throughout larval development, whereas the mNG equivalent lacked expression beyond the first larval stage (Fig. 3d to e). PEST also failed to destabilize nuclear or unlocalized mSG reporters driven throughout tissues using a heat shock promoter (Fig. 3f to i). Curiously, we observed distinct PEST activity with mNG::H2B reporters being effectively destabilized while unlocalized reporters were not effectively degraded, remaining detectable for 46 hours post-heat shock (Fig. 3f to i). This result was surprising as GFP::PEST reporters have been successfully used to capture oscillatory expression of numerous molting genes (Davis et al. 2004; Frand et al. 2005), and warrants further study. Together, our data argue that mSG is poorly degraded by a PEST sequence and therefore may be unsuitable as a reporter in most cases to capture dynamic gene expression. Rather, it is an outstanding lineage tracer for cells transiently expressing a gene of interest.

### Effect of epitope position and identity on PEST activity

The failure of our initial reporter to capture oscillatory *mlt-11* expression was due to the 3xFLAG tag N-terminal to the PEST sequence. This arrangement led to the most potent loss of PEST activity (Fig. 4a to c, 4j to l). The presence of 3xFLAG C-terminal or H2B sequences adjacent to PEST reduced PEST activity (Fig. 4h and i, 4m to o), albeit not to the same extent as the N-terminal 3xFLAG tag. Not all orientations of tag placement were tested in this study and there may be some instances where placing either of these tags next to a PEST sequence has minimal effect on oscillation. Unlike 3xFLAG, 3xMyc, 3xALFA, and 3xOLLAS had minimal impact on PEST-induced reporter destabilization (Fig. 5a to c, and e) when the tag was placed N-terminus to PEST. Expression levels, however, did change between these tags (Fig. 5f). 3xMyc was shown to produce higher reporter expression than 3xALFA, which in turn had higher expression than 3xOLLAS. These results indicate that while different epitopes may have an impact on PEST’s ability to destabilize a reporter, this does not necessarily translate to a loss of overall reporter oscillation. Curiously, 3xHA had a severe drop in expression and showed no significant oscillation (Fig. 3d and e). This result could reflect enhanced PEST activity or an inhibitory effect during transcription or translation. We favor the former explanation as HA tagging of proteins in cells has been performed routinely without widely reported issues with gene expression (Lo et al. 2013; Kuang et al. 2017; Toker et al. 2022).

PEST sequences are thought to promote ubiquitin-independent degradation through the 26S proteasome (Murakami et al. 1992). The 3xFLAG sequence could potentially disrupt this interaction, though why that would be specific to that epitope is unclear. 3xFLAG and 3xMyc are both negatively charged in the cell, with similar isoelectric points (pI=4.16 and 3.92 respectively) while a 3xHA tag has a similar pI (3.17) yet potentially increases PEST (pI=5.26) activity. 3xALFA and 3xOLLAS had no effect on PEST activity and are close to neutral (pI=6.27) or positively charged (pI=10.27) in cells, respectively. Notably, all of these epitopes were separated from the PEST sequence by a flexible 11 amino acid linker, so the effects of 3xFLAG and potentially that of 3xHA on PEST activity do not require being directly appended on the PEST sequence. While the mechanism behind disruption of PEST activity by N-terminal 3xFLAG epitopes is unclear, one should be aware when placing uncharacterized epitopes in close proximity to a PEST sequence.

### Best practices

Based on these results, it is recommended that, when using destabilized promoter reporters to assess dynamic gene expression, mSG be avoided. PEST::GFP::H2B or PEST::mNG::H2B performed comparably, and 3xFLAG epitopes adjacent to PEST should be avoided. The optimal configuration was PEST::[GFP or mNG]::H2B. If epitopes need to be used, 3xMyc, 3xALFA, and 3xOLLAS seem to be the safest options. In our hands, 3xOLLAS tags have outperformed 3xMyc tags for western blotting. We prefer nuclear localized promoter reporters as single-copy integrations were far easier to score compared to unlocalized reporters and in our heat shock assay unlocalized PEST::mNG::3xFLAG was not effectively destabilized (Fig 3g and i). mSG reporters appear to be better suited for lineage tracing and marking specific cell types. It will be important in the future to test whether the stabilizing effect of mSG on destabilized reporters is unique to PEST-based destabilization or whether it can stabilize other types of dynamically expressed proteins.

## METHODS AND MATERIALS

### Strains and Culture

*Caenorhabditis elegans* were cultured as described by (Brenner 1974), except worms were grown on MYOB media instead of NGM. MYOB agar was made as previously described by (Church et al. 1995). Animals were cultured at 20°C for all experiments and were grown at 15°C for general strain propagation. All created strains with genotypes can be found in Table 1.

**Table 1.** Strains Used in This Study.

| Name: | Genotype: |
| --- | --- |
| JDW719 | *jsSi1579 jsSi1706 jsSi1726 wrdSi116[loxP myo-2p::NLS::mNeonGreen, rps-0p HygR, loxP mlt-11p (-1977-5300 bp)::mNeonGreen (dpi)::3xFLAG::PEST::tbb-2 3'UTR FRT3] II* |
| JDW813 | *jsSi1579 jsSi1706 jsSi1726 wrdSi121[loxP myo-2p::NLS::mNeonGreen, rps-0p HygR, loxP mlt-11p (-1977-5300 bp)::PEST::GFP::H2B (his-58)::unc-54 3'UTR FRT3] II* |
| JDW837 | *jsSi1579 jsSi1706 jsSi1726 wrdSi126[loxP myo-2p::NLS::mNeonGreen, rps-0p HygR, loxP qua-1p (-2kb)::PEST::GFP::H2B (his-58)::unc-54 3'UTR FRT3] II* |
| JDW858 | *jsSi1579 jsSi1706 jsSi1726 wrdSi121[loxP myo-2p::NLS::mNeonGreen, rps-0p HygR, loxP mlt-11p (-1977-5300 bp)::GFP::H2B (his-58)::unc-54 3'UTR FRT3] II* |
| JDW859 | *jsSi1579 jsSi1706 jsSi1726 wrdSi121[loxP myo-2p::NLS::mNeonGreen, rps-0p HygR, loxP qua-1p (-2 kb)::PEST::mStayGold::H2B (his-58)::unc-54 3'UTR FRT3] II* |
| JDW861 | *jsSi1579 jsSi1706 jsSi1726 wrdSi121[loxP myo-2p::NLS::mNeonGreen, rps-0p HygR, loxP mlt-11p (-1977-5300 bp)::PEST::mStayGold::H2B (his-58)::unc-54 3'UTR FRT3] II* |
| JDW863 | *jsSi1579 jsSi1706 jsSi1726 wrdSi121[loxP myo-2p::NLS::mNeonGreen, rps-0p HygR, loxP mlt-11p (-1977-5300 bp)::PEST::mNG::H2B (his-58)::unc-54 3'UTR FRT3] II* |
| JDW865 | *jsSi1579 jsSi1706 jsSi1726 wrdSi128[loxP myo-2p::NLS::mNeonGreen, rps-0p HygR, loxP qua-1p (-2kb)::PEST::mNG::H2B (his-58)::unc-54 3'UTR FRT3] II* |
| JDW866 | *jsSi1579 jsSi1706 jsSi1726 wrdSi129[loxP myo-2p::NLS::mNeonGreen, rps-0p HygR, loxP qua-1p (-2kb)::PEST::mNG::3xFLAG::H2B (his-58)::unc-54 3'UTR FRT3] II* |
| JDW867 | *jsSi1579 jsSi1706 jsSi1726 wrdSi130[loxP myo-2p::NLS::mNeonGreen, rps-0p HygR, loxP qua-1p (-2kb)::GFP::H2B (his-58)::unc-54 3'UTR FRT3] II* |
| JDW869 | *jsSi1579 jsSi1706 jsSi1726 wrdSi131[loxP myo-2p::NLS::mNeonGreen, rps-0p HygR, loxP mlt-11p (-1977-5300 bp)::PEST::mNG::3xFLAG::H2B (his-58)::unc-54 3'UTR FRT3] II* |
| JDW895 | *jsSi1579 jsSi1706 jsSi1726 wrdSi131[loxP myo-2p::NLS::mNeonGreen, rps-0p HygR, loxP mlt-11p (-1977-5300 bp)::PEST::GFP::3xFLAG::H2B (his-58)::unc-54 3'UTR FRT3] II* |
| JDW901 | *jsSi1579 jsSi1706 jsSi1726 wrdSi131[loxP myo-2p::NLS::mNeonGreen, rps-0p HygR, loxP mlt-11p (-1977-5300 bp)::mNG::3xFLAG::PEST::H2B (his-58)::unc-54 3'UTR FRT3] II* |
| JDW911 | *jsSi1579 jsSi1706 jsSi1726 wrdSi131[loxP myo-2p::NLS::mNeonGreen, rps-0p HygR, loxP mlt-11p (-1977-5300 bp)::3xFLAG::PEST::GFP::H2B (his-58)::unc-54 3'UTR FRT3] II* |
| JDW912 | *jsSi1579 jsSi1706 jsSi1726 wrdSi131[loxP myo-2p::NLS::mNeonGreen, rps-0p HygR, loxP mlt-11p (-1977-5300 bp)::3xFLAG::PEST::mNG::H2B (his-58)::unc-54 3'UTR FRT3] II* |
| JDW932 | *jsSi1579 jsSi1706 jsSi1726 wrdSi137[loxP myo-2p::NLS::mNeonGreen, rps-0p HygR, loxP mlt-11p (-1977-5300 bp)::GFP::3xFLAG::PEST::H2B (his-58)::unc-54 3'UTR FRT3] II* |
| JDW943 | *jsSi1579 jsSi1706 jsSi1726 wrdSi137[loxP myo-2p::NLS::mNeonGreen, rps-0p HygR, loxP mlt-11p (-1977-5300 bp)::mNG::PEST::H2B (his-58)::unc-54 3'UTR FRT3] II* |
| JDW945 | *jsSi1579 jsSi1706 jsSi1726 wrdSi137[loxP myo-2p::NLS::mNeonGreen, rps-0p HygR, loxP mlt-11p (-1977-5300 bp)::H2B (his-58)::PEST::mNG::3xFLAG::unc-54 3'UTR FRT3] II* |
| JDW946 | *jsSi1579 jsSi1706 jsSi1726 wrdSi137[loxP myo-2p::NLS::mNeonGreen, rps-0p HygR, loxP mlt-11p (-1977-5300 bp)::mNG::PEST::3xFLAG::H2B (his-58)::unc-54 3'UTR FRT3] II* |
| JDW965 | *jsSi1579 jsSi1706 jsSi1726 wrdSi143[loxP myo-2p::NLS::mNeonGreen, rps-0p HygR, loxP hsp-16.48p::PEST::mNG::3xFLAG::unc-54 3'UTR FRT3] II* |
| JDW966 | *jsSi1579 jsSi1706 jsSi1726 wrdSi144[loxP myo-2p::NLS::mNeonGreen, rps-0p HygR, loxP hsp-16.48p::PEST::mStayGold::3xFLAG::unc-54 3'UTR FRT3] II* |
| JDW967 | *jsSi1579 jsSi1706 jsSi1726 wrdSi145[loxP myo-2p::NLS::mNeonGreen, rps-0p HygR, loxP hsp-16.48p::PEST::mNG::H2B (his-58)::unc-54 3'UTR FRT3] II* |
| JDW968 | *jsSi1579 jsSi1706 jsSi1726 wrdSi146[loxP myo-2p::NLS::mNeonGreen, rps-0p HygR, loxP mlt-11p::3xALFA::PEST::mNG::H2B (his-58)::unc-54 3'UTR FRT3] II* |
| JDW969 | *jsSi1579 jsSi1706 jsSi1726 wrdSi147[loxP myo-2p::NLS::mNeonGreen, rps-0p HygR, loxP mlt-11p (-1977-5300 bp)::3xMyc::PEST::mNG::H2B (his-58)::unc-54 3'UTR FRT3] II* |
| JDW973 | *jsSi1726 wrdSi150[loxP myo-2p::NLS::mNeonGreen, rps-0p HygR, loxP hsp-16.48p::PEST::mStayGold::H2B (his-58)::unc-54 3'UTR FRT3] II* |
| JDW974 | *jsSi1726 wrdSi151[loxP myo-2p::NLS::mNeonGreen, rps-0p HygR, loxP dpy-14p::PEST::mNG::H2B (his-58)::unc-54 3'UTR FRT3] II* |
| JDW975 | *jsSi1726 wrdSi152[loxP myo-2p::NLS::mNeonGreen, rps-0p HygR, loxP dpy-14p::PEST::mStayGold::H2B (his-58)::unc-54 3'UTR FRT3] II* |
| JDW796 | *jsSi1726 wrdSi153[loxP myo-2p::NLS::mNeonGreen, rps-0p HygR, loxP mlt-11p (-1977-5300 bp)::3xOLLAS::PEST::mNG::H2B (his-58)::unc-54 3'UTR FRT3] II* |
| JDW977 | *jsSi1726 wrdSi154[loxP myo-2p::NLS::mNeonGreen, rps-0p HygR, loxP mlt-11p (-1977-5300 bp)::3xHA::PEST::mNG::H2B (his-58)::unc-54 3'UTR FRT3] II* |

### Genome Editing

All plasmids used in this study can be found in Table 2. Annotated plasmid sequence files can be found in File S1. Specific cloning details and primers used are available upon request. In brief, a promoter and reporter would be integrated into a plasmid backbone via a SapTrap cloning reaction (Schwartz and Jorgensen 2016). This backbone would be capable of rapid recombination-mediated cassette exchange (rRMCE) at a specific locus of the *C. elegans* genome, allowing for genome editing to be consistent across all created strains (Nonet 2023).

**Table 2.** Plasmids Used in This Study.

| Name: | Construct: |
| --- | --- |
| pJW2361 | *mNeonGreen(dpi)::3xFLAG::PEST-tbb-2 3'UTR vector for SapTrap with ATG and GTA connectors* |
| pJW2387 | *Insertion vector for new hygR rapid RMCE approach* |
| pJW2408 | *hsp-16.48p with TGG and ATG connectors for SapTrap* |
| pJW2451 | *mlt-11p (Frand promoter -1977-5300 bp); SapTrap vector with TGG and ATG connectors.* |
| pJW2463 | *mlt-11p (-1977-5300 bp)::mNeonGreen (dpi)::3xFLAG::PEST::tbb-2 3'UTR in rapid RMCE vector* |
| pJW2640 | *PEST::GFP::H2B (his-58)::unc-54 3'UTR vector for rRMCE. Contains ClaI site for linearizing to Gibson clone in new promoters* |
| pJW2648 | *mlt-11p (-1977-5300 bp)::PEST::GFP::H2B (his-58)::unc-54 3'UTR vector for rRMCE* |
| pJW2650 | *qua-1p (-2 kb) SapTrap vector with TGG and ATG connectors.* |
| pJW2651 | *qua-1p (-2kb)::PEST::GFP::H2B (his-58)::unc-54 3'UTR vector for rRMCE* |
| pJW2698 | *PEST::mNG::H2B (his-58)::unc-54 3'UTR vector for SapTrap with ATG and GTA connectors* |
| pJW2699 | *PEST::mNG::3xFLAG::H2B (his-58)::unc-54 3'UTR vector for SapTrap with ATG and GTA connectors* |
| pJW2700 | *PEST::mStayGold::H2B (his-58)::unc-54 3'UTR vector for SapTrap with ATG and GTA connectors* |
| pJW2715 | *GFP::H2B (his-58)::unc-54 3'UTR vector for SapTrap with ATG and GTA connectors* |
| pJW2718 | *mlt-11p (-1977-5300 bp)::PEST::mNG::H2B (his-58)::unc-54 3'UTR vector for rRMCE* |
| pJW2719 | *mlt-11p (-1977-5300 bp)::PEST::mSG::H2B (his-58)::unc-54 3'UTR vector for rRMCE* |
| pJW2720 | *qua-1p (-2kb)::PEST::mNG::H2B (his-58)::unc-54 3'UTR vector for rRMCE* |
| pJW2721 | *qua-1p (-2kb)::PEST::mSG::H2B (his-58)::unc-54 3'UTR vector for rRMCE* |
| pJW2722 | *mlt-11p (-1977-5300 bp)::PEST::mNG::3xFLAG::H2B (his-58)::unc-54 3'UTR vector for rRMCE* |
| pJW2724 | *qua-1p (-2kb)::PEST::mNG::3xFLAG::H2B (his-58)::unc-54 3'UTR vector for rRMCE* |
| pJW2726 | *mlt-11p (-1977-5300 bp)::GFP::H2B (his-58)::unc-54 3'UTR vector for rRMCE* |
| pJW2727 | *qua-1p (-2kb)::GFP::H2B (his-58)::unc-54 3'UTR vector for rRMCE* |
| pJW2729 | *PEST::GFP::3xFLAG::H2B (his-58)::unc-54 3'UTR vector for SapTrap with ATG and GTA connectors* |
| pJW2730 | *PEST::mNG::3xFLAG::unc-54 3'UTR vector for SapTrap with ATG and GTA connectors* |
| pJW2731 | *PEST::mStayGold::3xFLAG::unc-54 3'UTR vector for SapTrap with ATG and GTA connectors* |
| pJW2767 | *mlt-11p (-1977-5300 bp)::PEST::GFP::3xFLAG::H2B (his-58)::unc-54 3'UTR vector for rRMCE* |
| pJW2779 | *mNG::3xFLAG::PEST::H2B (his-58)::unc-54 3'UTR vector for SapTrap with ATG and GTA connectors* |
| pJW2781 | *3xFLAG::PEST::GFP::H2B (his-58)::unc-54 3'UTR vector for SapTrap with ATG and GTA connectors* |
| pJW2782 | *3xFLAG::PEST::mNG::H2B (his-58)::unc-54 3'UTR vector for SapTrap with ATG and GTA connectors* |
| pJW2789 | *mlt-11p (-1977-5300 bp)::mNG::3xFLAG::PEST::H2B (his-58)::unc-54 3'UTR vector for rRMCE* |
| pJW2808 | *mlt-11p (-1977-5300 bp)::3xFLAG::PEST::GFP::H2B (his-58)::unc-54 3'UTR vector for rRMCE* |
| pJW2809 | *mlt-11p (-1977-5300 bp)::3xFLAG::PEST::mNG::H2B (his-58)::unc-54 3'UTR vector for rRMCE* |
| pJW2850 | *GFP::3xFLAG::PEST::H2B (his-58)::unc-54 3'UTR vector for SapTrap with ATG and GTA connectors* |
| pJW2851 | *mNG::PEST::H2B (his-58)::unc-54 3'UTR vector for SapTrap with ATG and GTA connectors* |
| pJW2853 | *H2B::PEST::mNG::3xFLAG::unc-54 3'UTR vector for SapTrap with ATG and GTA connectors* |
| pJW2856 | *mNG::PEST::3xFLAG::H2B (his-58)::unc-54 3'UTR vector for SapTrap with ATG and GTA connectors* |
| pJW2887 | *mlt-11p (-1977-5300 bp)::GFP::3xFLAG::PEST::H2B (his-58)::unc-54 3'UTR vector for rRMCE* |
| pJW2897 | *mlt-11p (-1977-5300 bp)::mNG::PEST::H2B (his-58)::unc-54 3'UTR vector for rRMCE* |
| pJW2899 | *mlt-11p::H2B (his-58)::PEST::mNG::3xFLAG::unc-54 3'UTR vector for rRMCE* |
| pJW2900 | *mlt-11p (-1977-5300 bp)::mNG::PEST::3xFLAG::H2B (his-58)::unc-54 3'UTR vector for rRMCE* |
| pJW2916 | *3xOLLAS::PEST::mNG::H2B (his-58)::unc-54 3'UTR vector for SapTrap with ATG and GTA connectors* |
| pJW2917 | *3xALFA::PEST::mNG::H2B (his-58)::unc-54 3'UTR vector for SapTrap with ATG and GTA connectors* |
| pJW2918 | *3xHA::PEST::mNG::H2B (his-58)::unc-54 3'UTR vector for SapTrap with ATG and GTA connectors* |
| pJW2919 | *3xMyc::PEST::mNG::H2B (his-58)::unc-54 3'UTR vector for SapTrap with ATG and GTA connectors* |
| pJW2920 | *dpy-14p (-279 bp) SapTrap vector with TGG and ATG connectors.* |
| pJW2921 | *hsp-16.48p::PEST::mNG::3xFLAG::unc-54 3'UTR vector for rRMCE* |
| pJW2922 | *hsp-16.48p::PEST::mStayGold::3xFLAG::unc-54 3'UTR vector for rRMCE* |
| pJW2928 | *hsp-16.48p::PEST::mNG::H2B::unc-54 3'UTR vector for rRMCE* |
| pJW2929 | *hsp-16.48p::PEST::mStayGold::H2B::unc-54 3'UTR vector for rRMCE* |
| pJW2930 | *mlt-11p (-1977-5300 bp)::3xALFA::PEST::mNG::H2B (his-58)::unc-54 3'UTR vector for rRMCE* |
| pJW2931 | *mlt-11p (-1977-5300 bp)::3xMyc::PEST::mNG::H2B (his-58)::unc-54 3'UTR vector for rRMCE* |
| pJW2932 | *dpy-14p::PEST::mNG::H2B::unc-54 3'UTR vector for rRMCE* |
| pJW2933 | *dpy-14p::PEST::mStayGold::H2B::unc-54 3'UTR vector for rRMCE* |
| pJW2934 | *mlt-11p (-1977-5300 bp)::3xOLLAS::PEST::mNG::H2B (his-58)::unc-54 3'UTR vector for rRMCE* |
| pJW2935 | *mlt-11p (-1977-5300 bp)::3xHA::PEST::mNG::H2B (his-58)::unc-54 3'UTR vector for rRMCE* |

### Imaging Worms

Synchronized worms were gathered by picking directly off of MYOB plates. Worms would be transferred to 24µl M9 + 5 mM levamisole on a 2% agarose pad on a slide and secured with a coverslip. Images were acquired using a Plan-Apochromat 40x/1.3 Oil DIC lens or a Plan-Apochromat 63x/1.4 Oil DIC lens on an AxioImager M2 microscope (Carl Zeiss Microscopy, LLC) equipped with a Colibri 7 LED light source and an Axiocam 506 mono camera. Acquired images were processed through Fiji software (version: 2.16.0/1.54p). Within each figure, unless noted elsewhere, the same exposure times (250 ms) were used for DIC and 488 nm excitation/green emission (AF488) channel imaging. Figure 1 used 1.75 s for imaging mNeonGreen. All experiments used an exposure time to avoid saturation of the brightest samples, allowing for equal comparison between experimental groups. For the heat shock experiments, L4 worms were picked to a fresh MYOB plate and incubated at 34°C for 1 hour to induce expression of the *hsp-16*.*48* promoter reporters. Worms were immediately imaged at 100X magnification with an exposure time of 50 ms for expression in the cytoplasm and 150 ms for nuclear expression. Heat shocked worms were maintained at room temperature for 3, 24 and 46 hours and then imaged at these time points. The area immediately adjacent to the vulva and intestinal nuclei were chosen for cytoplasmic and nuclear imaging, respectively, due to their uniform size and structure throughout development.

### Scoring reporter expression

Animals were synchronized by timed egg lay, wherein worms were picked to a new plate, allowed to lay eggs for 1 hour, then removed from the plate. The 0 hour timepoint would then be set to the moment the P0 animals were removed. Animals were scored for reporter expression using a PlanApo 5.0×/0.5 objective on a M165 FC stereomicroscope (Leica) equipped with an X-cite FIRE LED lightsource (Excelitas) and long-pass GFP filter set (Leica, 10447407) set to differing LED light intensities. In experiments that use differing percentages for fluorescence, each animal would first be scored with the LED light intensity set to 33%, then to 100%. This allowed the scorer to more easily detect when the promoter reporter was on and to what degree. In experiments that scored animals over a 8 hour period, 10% LED light intensity was used as the scorer could reliably detect expression. For each subfigure, 25 animals were scored over two experiments for a total of 50 animals. The promoter reporter was only considered on if the green H2B signal was clearly visible in the hypodermis, and any diffuse green was considered off for consistency.

### Mean Intensity Value Measurements

Worms were imaged over two experiments. Each worm had its five brightest nuclei measured by a circle using Zen 3.4 (Zen lite) of area 10.012 µm^2^ and a diameter of 3.570 µm. We measured the mean intensity value of each circle. The five nuclei were averaged per worm and plotted in GraphPad Prism 10 to compare data and run statistical analysis. For heat shock experiments, nuclear reporters measured one intestinal nucleus per worm with a 100 µm^2^ box. For cytoplasmic reporters, one box was drawn for each worm adjacent to the vulva where expression was consistent throughout development. Details of statistical tests that we performed can be found in the figure legends where applicable.

## Data availability

A full description of all plasmids and transgenic *C. elegans* are in Tables 1 and 2. The sequence of the plasmids used to generate transgenic strains is in File S1 which will be available through FigShare. The authors affirm that all data necessary for confirming the conclusions of the article are present within the article, figures, tables, and supplementary material. Any additional information, such as oligonucleotide sequences or cloning approaches will be provided upon request.

## Acknowledgements

The authors thank Tabatha Wells, McKenna Berry, and Janet Noriega for research support. Some strains were provided by the Caenorhabditis Genetics Center, which is funded by the NIH Office of Research Infrastructure Programs (P40 OD010440). Wormbase was used in the design and execution of experiments.

